# A framework for space-efficient variable-order Markov models

**DOI:** 10.1101/443101

**Authors:** Fabio Cunial, Jarno Alanko, Djamal Belazzougui

## Abstract

**Motivation:** Markov models with contexts of variable length are widely used in bioinformatics for representing sets of sequences with similar biological properties. When models contain many long contexts, existing implementations are either unable to handle genome-scale training datasets within typical memory budgets, or they are optimized for specific model variants and are thus inflexible.

**Results:** We provide practical, versatile representations of variable-order Markov models and of interpolated Markov models, that support a large number of context-selection criteria, scoring functions, probability smoothing methods, and interpolations, and that take up to 4 times less space than previous implementations based on the suffix array, regardless of the number and length of contexts, and up to 10 times less space than previous trie-based representations, or more, while matching the size of related, state-of-the-art data structures from Natural Language Processing. We describe how to further compress our indexes to a quantity related to the redundancy of the training data, saving up to 90% of their space on repetitive datasets, and making them become up to 60 times smaller than previous implementations based on the suffix array. Finally, we show how to exploit constraints on the length and frequency of contexts to further shrink our compressed indexes to half of their size or more, achieving data structures that are 100 times smaller than previous implementations based on the suffix array, or more. This allows variable-order Markov models to be trained on bigger datasets and with longer contexts on the same hardware, thus possibly enabling new applications.

**Availability and implementation:** https://github.com/jnalanko/VOMM

## 1 Introduction

Building statistical models for large sets of sequences that share biological properties is a fundamental problem in bioinformatics, with connections to coding, compression, and machine learning. In many biological sequences, the empirical probability distribution of the next character does not change significantly if one takes into account a subsequence of the recent history (called the *context* of the character) that is longer than a fixed threshold, thus Markov models in which all states are strings of uniform length k, called the *order* of the model, are routinely used. Choosing the order of a Markov model asks one to balance between conflicting issues. Longer contexts imply estimating exponentially more parameters, which might not be feasible with the available data and which might hurt generalization. However, while most long contexts might be too infrequent to yield accurate probability estimates, a few of them might still occur frequently enough to be used as predictors in practice. Shorter contexts, on the other hand, yield models of tractable size, but they might lead to inaccurate predictions, since they are unable to capture long-range dependencies. *Variable-order Markov models* (VOMMs, [42]) aim at solving such issues by allowing the length of the context used to predict the next character to depend on the recently observed characters themselves. This dependency, called *local order*, has indeed been observed in a number of natural sources [8]. A VOMM can thus be seen as a compression of a fixed-order Markov model of large order, in which all states that yield similar character distributions are collapsed into a single state, whose sequence is decided by a number of statistical criteria.

Variable-order Markov models and their variants, like *interpolated Markov models* (IMMs, see e.g. [46]), are by now a staple of bioinformatics, and have been successfully applied to detect domains in protein sequences [4], to segment protein sequences into domains [50, 8, 6], to assign domains and proteins to families [4], to separate coding from non-coding DNA regions [45], to detect horizontal gene transfer [21], to identify genes in newly sequenced microbial genomes after being trained on known open reading frames [46, 24, 25], and to detect eukaryotic promoters [39]. VOMMs and their variants have been applied to metagenomic samples as well. For example, VOMMs have been used to separate the reads of a eukaryotic host from those of an intracellular prokaryotic parasite [24]; to model known genomes in order to estimate, given a metagenomic sample, the genome or taxon a read was sampled from [12]; to define compositional distances between metatranscriptomic samples [32]; and to model the clusters produced by reference-free binning of metagenomic reads [28, 56, 58].

As mentioned, a key reason for the introduction of VOMMs is the implicit compression they perform over a Markov model of fixed, large order [42, 62]. Space-efficient implementations of VOMMs and of their variants are becoming increasingly relevant in the post-genome era, in which one wishes to train them on long genomes, on large metagenomic samples, or on massive collections of genomes of similar species or of proteins with the same function or structure [6, 7, 8, 28]. When the criteria for deciding whether a subsequence of the training data should be used as a context are so stringent that just a small set of short subsequences is chosen, the space taken by a VOMM is not a practical issue, and a simple pointer-based trie implementation like e.g. the one described by [5] suffices. In this paper we focus on VOMMs in which the total length of all contexts is large enough to make such a simple approach impractical. This is a concrete scenario, especially in large datasets, since e.g. increasing the maximum length of a context does not decrease classification performance in practice [4], and in real use cases one wishes to experiment with multiple selection criteria without worrying about blowups in space or training time.

The first training algorithms required *O*(*k|T |*^2^) time for building a VOMM from a sequence *T*, where *k* is an upper bound on context length, and *O*(*|S|*^2^) time to compute the probability of a query sequence *S* according to the model [45]. Later, [1] described a general-purpose data structure, based on the suffix tree of T, that can be built in *O*(*|T |*) time for alphabets of constant size σ and for a large number of context selection criteria, and that allows one to score a query *S* in *O*(*|S|*) time for constant alphabets. Such data structure was the first to take *O*(*|T |* log *|T |*) bits of space, regardless of the number and length of contexts, and it was later implemented using lazy suffix trees or enhanced suffix arrays [48, 49]. However, such implementations are not available any more [47]. [33] designed and implemented another data structure, based on the suffix array and the inverse suffix array of *T*, that takes again *O*(*|T |* log *|T |*) bits of space, but that allows scoring a query using as context just the longest match with the training data at each position. Meanwhile, IMMs with ad hoc emission probability formulas were being deployed in natural language processing (NLP), and compact indexes for computing a specific family of such probabilities have been developed and implemented at scale by [52, 53]. Such data structures rely on compressed suffix trees, and thus need just *O*(*|T |* log σ) bits of space. However, they do not implement general purpose VOMMs, and they are optimized to answer queries on the probability of observing a character after an arbitrarily large number *ℓ*of other characters specified in the query, rather than to score an entire sequence *S*, thus they take *O*(*ℓ|S|*) time to compute such global score in the worst case (although practical implementations try to reuse counts across adjacent sliding windows). More recently, [10] described a general-purpose data structure that takes just 3*|T |* log σ + *o*(*|T |* log σ) bits of space to implement a VOMM or an IMM, regardless of the number and length of contexts and for a large set of context selection criteria, and that can be trained in *O*(*|T |*) time using *O*(*|T |* log σ) bits of working space. In theory the space taken by such data structure is similar to the space taken by the data structures by [52, 53], while at the same time supporting a variety of context selection and probability smoothing criteria, and allowing one to score a query *S* in *O*(*|S|*) time using 2*|S|* + *o*(*|S|*) bits of space.

This paper is concerned with building *practical, general-purpose representations of VOMMs and IMMs that are as small as possible*. Optimizing query time and the construction of such representations is outside the scope of the paper. Our first contribution is an implementation of the theory described in [10]: we provide a general-purpose framework of practical data structures and algorithms that allows bioinformaticians to implement a large number of VOMMs and IMMs in small space, including many context-selection criteria, scoring functions, probability smoothing methods, and interpolations, without affecting the accuracy of the models and scaling to large datasets. No such framework currently exists, either because available implementations cannot handle large datasets within typical memory budgets, or because they support just few specific Markov models. We implement a representative subset of all the variants that our framework supports, achieving up to approximately 4 times less space than [33] and 10 times less space (or even less, depending on context selection criterion) than [5], while matching the space of the NLP data structures in [52, 53]. Since VOMMs are generative statistical models, their training datasets consist typically of positive examples of a class, and are thus likely to be repetitive. The second contribution of this paper is an extension of the approach in [10] which, for the first time, reduces the size of a VOMM to a quantity related to the redundancy of the training data. In repetitive datasets, this allows saving up to approximately 90% of the space of our index, without any effect on accuracy, making our data structures become up to approximately 60 times smaller than [33]. Finally, we describe a way of shrinking our data structures even further when the user sets an upper bound on context length, or a lower bound on context frequency, which are common in applications (see e.g. [18, 45, 46]). This can shrink our already compressed index to half of its size and beyond, depending on the dataset, yielding a data structure that is up 100 times or more smaller than [33], again without affecting accuracy. Such space reductions apply to a large number of VOMM and IMM variants, albeit not to all of them, and, as expected, they come at the cost of increased scoring time: even the basic version of our index is between 35 and 60 times slower than [5], and between 3 and 12 times slower than [33].

## 2 Preliminaries and notation

### 2.1 Strings and string indexes

Let Σ= [1..σ] be an integer alphabet, let # = 0 be a separator not in Σ, and let *S* and *T* be strings in [1..σ]*. The *matching statistics array* MS_*S,T*_ is an array of length *|S|* such that MS_*S,T*_ [*i*] is the largest *ℓ* such that *S*[*i*–*ℓ*+ 1..*i*] occurs in *T*. We denote by W̅ the reverse of a string *W*, i.e. string *W* read from right to left. We denote by *f_T_*(*W*)the number of (possibly overlapping) occurrences of a string *W* in *T*, we denote by 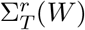 the set of distinct characters that occur to the right of *W* in *T*, and we denote by 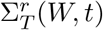 (respectively, by 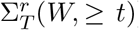 the set of distinct characters that occur exactly (respectively, at least) *t* times to the right of *W* in *T*. The characters in 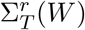 are called *right-extensions* of *W*, and 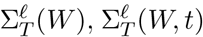 and 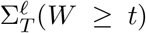 are defined symmetrically for the left side of *W*. We use small sigma as a short-hand for the cardinality of every such set, e.g. 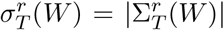. We call *empirical emission probability* of a character a from a substring *W* of *T* the quantity *P_T_* (*a|W*) = *f*_T_ (*Wa*)/*f*_T_ (*W*), although a number of other estimates have been proposed, for example the modified Laplace rule *P*(*a|W*) = (*f*(*W a*) + 0.5)/(*f* (*W*) + *σ*/2) and *P*(*a|W*) = *f*(*W a*)/(*f*(*W*) + *σ*^r^(*W*)) (see e.g. [1, 3, 62, 64, 40, 8, 4, 2, 18]). We call *empirical probability of a string W* in *T* the quantity *P_T_* (*W*) = *f_T_* (*W*)/(*|T | – |W |* + 1). A *repeat W* of *T* is a substring of *T* that satisfies *f*(*W*) > 1. *W* is *right-maximal* (respectively, *left-maximal*) iff 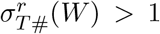 (respectively, iff 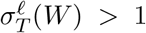 We call *right-deterministic* (respectively, *left-deterministic*) a string that is not right-maximal (respectively, left-maximal). It is well known that *T* can have at most *|T |-*1 right-maximal repeats and at most *|T |-*1 left-maximal repeats. A *maximal repeat* is a repeat that is both left- and right-maximal.

We assume the reader to be familiar with standard notions from text indexes, including the suffix tree ST_*T*_ and the suffix-link tree SLT_*T*_ of a string *T*, Weiner links, the Burrows-Wheeler transform BWT_*T*_, the lexicographic interval of a right-maximal repeat in the BWT, backward steps, rank and select queries, wavelet trees, and compact representations of tree topologies. See Section 9 in the supplement for more details and for pointers to the literature. In the rest of the paper, we omit subscripts whenever they are clear from the context.

### 2.2 Variable-order Markov models

We assume the reader to be familiar with fixed-order Markov models, which we do not define or further describe here. Given a Markov model of fixed order *k* on alphabet [1..σ], let ℙ (*a|X*) be the *emission probability* of state *X*, i.e. the probability of observing character a after the length-k context X. Given a string W of length *h* < *k*, it might happen that ℙ (*a|UW*) = ℙ (*a|V W*) for all strings *U* and *V* of length *k – h* and for all characters *a*: in other words, adding up to *k – h* characters before *W* does not alter the probability of seeing any character after *W*. This is called *variable-order Markov model* (VOMM), and it can be represented more compactly by storing just *W* (rather than all *k*-mers with *W* as a suffix) and the emission probabilities of *W* [42]. In what follows we assume that a Markov model is built from a training string *T*. The set of contexts is typically stored in a trie 𝒯, called *context tree* or *probabilistic suffix tree* (see e.g. [23, 7] and references therein), where contexts are inserted from right to left, and in which every node that corresponds to a context is marked and stores emission probabilities [45]. We denote by *𝒞* the set of nodes of *𝒯* that are marked as contexts (abusing notation, we use symbol *𝒞* to denote also the set of contexts).

In the following sections we survey a number of criteria used to assign a probability to a query, and to select contexts from substrings of the training data. As we will see in Section 4, most such criteria can be implemented with a small setup of space-efficient data structures.

#### 2.2.1 Scoring a query

The probability of a query string *S* according to the VOMM is computed character by character, as follows: at every position *j* of *S*, the probability of character *S*[*j*] = *b* is decided by finding a context *w* ∈ 𝒞 based on *S*[1.. *j –*1], and by accessing the empirical probability *P*(*b|W*) stored at *w* [43, 42]. The function that returns the context to be used at position *j* is called *structure function* [43], and it typically returns the longest context that is a suffix of *S*[1..*j –* 1]. The probability of *S* is then the product of the probability of every character of *S* (in the rest of the paper we focus on computing log-probabilities, thus all products become additions). Some variants multiply the probability of *S* by the average value of σ^r^(*W*) over all contexts *W* used for prediction [37].

In most applications, when *W* does not emit *b* in the model, the probability of character *S*[*j*] should still be set to a nonzero value: deciding such nonzero value is called the *zero-frequency* or the *sparse data problem* [18, 64, 4], and taking this into account to adapt the probability of characters that are emitted by the model is called *smoothing*. Such nonzero value can be a user-defined constant, a function of *f*(*b*), *f*(*W*), σ, σ^r^(*W*) or σ^*r*^(*W*, 1), like the Good-Turing estimator (see e.g. [42, 4, 18, 17]), or it can be computed recursively. For example, in the conceptually similar PPM compression algorithm [18], one assigns to *S*[*j*] a probability *P*̃(*b|W*) defined as follows [3]: *P*̃(*b|W*) = *P*(*b|W*) *·* (1 *– P̂(W)·*if *W* ∈ 𝒞 and *P̃*(*b|W)>0, P*̃(*b|W*) = *P̂(W)· P*̃(*b|W* [2..*|W |*]) otherwise, where *P̂* (*W*), called *escape* or *backoff probability*, is an estimate of the probability of observing a character that does not follow *W* in the model, for example the Good-Turing estimator mentioned above (see e.g. [3, 18, 64] and references therein). In some approaches, recursion stops when the suffix reaches a user-specified minimum length [39, 46, 13] or frequency [46].

A similar approach can be applied even when W emits b, to compute a *mixture* of probability estimates (see e.g. [16, 46]), possibly with weights that depend on the length of the contexts [23]: this is also called *blending* [35]. An *interpolated Markov model* (IMM) is a fixed-order Markov model in which emission probabilities are a weighted mixture of lower-order contexts [46, 39], for example p̃(*b|W*) =, λ(*W*) *P*(*b*|*W*)+(1– λ(*W*)). p̃(*b| W*[2..|W|]) if |*W*|>0, and otherwise it is a function of quantities like *f*(*b*),|*T*|, σ and σ^*r*^(*W*) (see e.g.[16]). Weights λ(*W*) can represent the confidence in the accuracy of an emission probability estimate (longer contexts yield stronger predictions, but shorter contexts have more accurate statistics), or the prior that correlation decreases with distance[23,22,17]. Weights can be a fixed vector of constants (which might sum to one), or a given function of the frequency, length, and number of right-extensions of every suffix (see e.g. [39, 60, 46, 44, 16]). More complex variants of recursive emission probability functions have been designed in natural language modeling and information retrieval, e.g. Kneser-Ney smoothings (see e.g. [52, 53, 51, 30] and references therein). Non-recursive ways of interpolating the probabilities of different contexts have also been proposed. For example, one can set *P*̃(*a|W*) = (*f*(*W a*) *– x*)/*f*(*W*) +*α*(*W*)*β*(*W*, *α*) if *f*(*W* α) > 0, and *P*̃(*a|W*) = α(*W*)*β*(*W*, *a*) otherwise, where *x* is a parameter such that *x* < *f*(*Wc*) for all *c* ∈ Σ^*r*^ (*W*), *β*(*W*, *a*) = σ^ℓ^(*W* [2..*|W |*]*a*)/*f*(*W* [2..*|W |*]) is a back-off function, and α(*W*) = σ^*r*^(*W*) *x*/*f*(*W*) [29]. Yet other variants take the *maximum* emission probability over all possible contexts, rather than blending context probabilities together [37].

#### 2.2.2 Selecting contexts

A number of algorithms for learning context trees optimally in a statistical sense have been described (see e.g. [22, 63, 45, 27, 15], including methods that exploit connections with the suffix tree and BWT [2, 35]. Here we focus on criteria that select a substring *W* of the training data as a context based only on properties of *W*. *Combinatorial approaches* mark all sub-strings as contexts [33] and select the longest context at each position of the query, or alternatively mark all right-deterministic substrings as contexts [59, 19], and start prediction from the shortest context at each position of the query (if any). *Statistical approaches* mark as contexts all the frequent substrings *aW* (where *a* ∈ [1..σ] and *W* ∈ [1..σ]^*^) having at least one character b with high emission probability from *aW*, and such that the emission probability of *b* from *aW* is significantly different from the emission probability of b from W (see e.g. [7] and references therein). Specifically, for user-defined thresholds τ_1_ > 0, τ _2_ > 0, τ _3_ < 1 and τ _4_ > 1, they require *P*(*aW*) ≥ τ _1_, *P*(*b|aW*) ≥ τ _2_, and *P*(*b|aW*)/*P* (*b|W*) *∈* (0.. τ _3_] ∪ [τ _4_.. + *∞*). Alternatively, contexts are all the substrings *aW* of *T* with high Kullback-Leibler divergence between the probability distribution of the characters that follow *aW*, and the probability distribution of the characters that follow *W* [15, 62, 14], i.e. ∑_b∈[1..σ]_ *f*(*aW b*) log(*P*(*b|aW*)/*P* (*b|W*)) ≥ τ for a positive τ. KL divergence is sometimes replaced by the squared *L*_1_ norm [15], or by any *p*-norm, again using a positive threshold. In yet other variants (e.g. [42, 61, 62]), a substring *W* of *T* is called *unstable* and it is used as a context if it has high entropy compared to all its left-extensions, i.e. if *f*(*W*)*H*(*W*) *–*∑_a∈[1..σ]_ *f*(*aW*)*H*(*aW*) ≥ τ, where the entropy *H*(*W*)equals–∑_b∈[1..σ]_*P*(*b*|*W*) log *P*(*b*|*W*) and τ is a positive threshold. This can be extended to longer left-extensions of a user-specified length. Alternatively, one can use the shortest *stable* suffix of the current history as a context [61].

### 2.3 Variable-order Markov models in small space

To keep the paper self-contained, we summarize here the key ideas of [10, Section 6]. It can been shown that the statistical criteria for choosing contexts in Section 2.2.2 can only select either a *maximal repeat*, or the *left-extension by one character of a maximal repeat* (including possibly strings that occur just once in *T*). In Section 1 of the supplement we show that such property holds also for a class of IMMs that have been widely used in gene identification [46]. The algorithm in [10] supports any selection criterion in which contexts are maximal repeats or left-extensions thereof: the context tree of *T* is then the sub-graph of the extended suffix-link tree 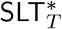, defined by the following procedure: starting from every node of 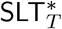 that corresponds to a context, recursively follow reverse Weiner links up to the root of 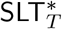, marking all nodes met in the process.

Our index on *T* consists of 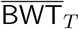 and the topologies of 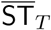 and SLT_T_. We represent 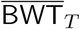 as a wavelet tree that supports Weiner links in *O*(log σ) time, and we use standard compact data structures to support constant-time queries on the balanced parentheses representations of 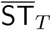 and SLT_T_ (see Section 9 in the supplement). Let `idSLT`(*υ*) be the position of node υ of SLT in the preorder traversal of SLT. We build a bitvector `mrSLT`[1..*p*], with a bit associated to each of the *p* nodes of SLT, such that `mrSLT`[`idSLT`(v)] = 1 iff υ is a maximal repeat. Similarly, let `idST`(υ) be the position of node υ of 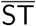 in the preorder traversal of 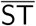. We build another bitvector `mrST`[1..*q*], with a bit for each of the *q* nodes of 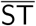, such that `mrST`[`idST`(υ)]=1 if υ is a maximal repeat. We index `mrSLT` to support select queries, and `mrST` to support rank and select queries. Since SLT is a subdivision of the subgraph of 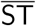 induced by maximal repeats, the *i*-th one in `mrST` and the *i*-th one in `mrST` correspond to the same maximal repeat. Thus, if node υ is a maximal repeat and if we know `idST`(υ), we can compute the length of *ℓ*(υ) by going to the node υ′ in SLT with `idSLT`(υ′) = select(`mrSLT`, 1, `rank`(`mrST`, 1, `idST`(υ))) and by computing the depth of υ′ in the topology of SLT. In other words, SLT can be seen as a data structure that stores the lengths of all maximal repeats as tree depths. Finally, we include in the index a bitvector context[1..p] such that context[`idST`(υ)] = 1 iff node υ in 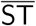 is the locus of a context, and a lowest marked ancestor data structure [57] to move in constant time from any node in the topology of 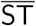 to its lowest ancestor that is as context.

We summarize how to implement just one of the scoring methods of Section 2.2.1, namely, starting from the longest context *W* that ends at position *i* of the query *S*, and using the PPM recursive scoring *P*̃(*S*[*i*+1]*|W*) = *P*(*S*[i+1]*|W*) *·* (1–*P*̂ (*W*)) if *W* is a context that emits *S*[*i* + 1], and otherwise *P*̃ (*S*[*i* + 1]*|*i) = *P*̂ (*W*) *· P*̃ (*S*[*i* + 1]*|W* [2..*|W |*]), where *P*̂ (*W*) is either a constant, or a function of *f*(*S*[*i* + 1]), *f*(*W*), σ, *σ^r^* (*W*) or *σ^r^* (*W*, 1). Recursion stops at the longest context that ends at position *i* and emits *S*[*i*+1]. Section 4 details other variants.

The algorithm maps the process of scoring every position *i* of *S* onto computing the matching statistics value **MS**[*i*], keeping the invariant of knowing, for each *i*, **MS**[*i*] itself, and the locus of the reverse of *S*[*i -* **MS**[*i*] + 1..*i*] in 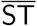. The actual **MS** array is never fully present in memory, since **MS**[*i*] values are computed on the fly. Assume that contexts are a subset of the maximal repeats of *T*. We scan S from left to right, issuing Weiner link and parent operations on 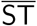 as done by standard matching statistics algorithms (see e.g. [38, 9]): such operations can be implemented using just 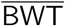 and the topology of 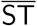.

Assume that, after taking a sequence of successful Weiner links from the first position of *S*, we are currently at position *i* in *S*. Using the topology of 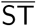, we convert the interval of the reverse of *W* = *S*[1..*i*] in 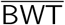 into the `idST` value of its locus υ in 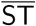. Note that the label of υ is either the reverse of *W*, or the reverse of *XW*, where *X* is a nonempty string. We then check whether context[`idST`(υ)] = 1. If this is true, then we can measure *|ℓ*(υ)*|* in constant time since υ is a maximal repeat, and if it is equal to *i* we know that *W* is the longest context that ends at position i. If *|ℓ*(υ)*|* > *i*, or if context[`idST`(υ)] ≠1,we move in constant time to the lowest ancestor υ′ of υ that is marked as a context: its label is clearly the longest context that ends at *i*, and the length of its label is necessarily smaller than *i*. We compute the emission probability of character *S*[*i* + 1] by taking the Weiner link with character *S*[*i* + 1] from υ′. If such Weiner link exists, or if no such Weiner link exists and we choose not to back off to shorter contexts, we are done. Otherwise, we iteratively issue parent operations from υ′, checking whether the nodes we meet are contexts and if they have a Weiner link by character *S*[*i* + 1]. If this is the case, recursion stops. Otherwise, it continues, and since υ′ is a context, all the nodes we meet are maximal repeats and we can measure the length of their labels, thereby implementing the backoff formula. Once we have assigned a score to position *i* + 1 of *S*, we continue as in a standard matching statistics algorithm: we keep increasing *i* and issuing a Weiner link with character *S*[*i* + 1] from the locus of the reverse of *S*[1..*i*] in 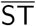. When a Weiner link fails with character *S*[*i* + 1], we keep issuing parent operations on 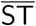 until we reach a node with a Weiner link labeled by *S*[*i* + 1]: the label of such node is a maximal repeat of *T*, its length is MS[*i*], and we can measure such length using SLT: we have thus re-established the invariant for the new value of *i*, and we can repeat the whole process. Just minor modifications are required when contexts are left-extensions by one character of maximal repeats: see Section 2 in the supplement for details. It can be shown that all the operations of the algorithm, including those in the recursive backoff, map onto operations performed by a matching statistics algorithm, and this number is *O*(*|S|*); each operation takes *O*(log σ) time, since Weiner links are the bottleneck.

## 3 Smaller variable-order Markov models

Recall that the algorithm in Section 2.3 works for contexts that are maximal repeats or left-extensions thereof. In repetitive strings, the number of maximal repeats can be significantly smaller than the number of suffix tree nodes [11], thus one could try to shrink the size of the data structures to a quantity related to the number of maximal repeats of the training data. We first show how to do this for computing MS_*S,T*_:

### Lemma 1

MS_*S,T*_ *can be computed using data structures of size proportional to the number of left extensions of maximal repeats of T.*

*Proof.* Let *V* be the set of nodes of 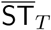, let *M* ⊂*V* be the set of nodes that correspond to maximal repeats, and let *L* ⊂*V* be the set of leaves. We replace the topology of 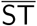 with the topology of the subgraph *G* of 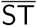 induced by M∪ L, then we collapse into the same leaf every maximal interval of leaves of G that are consecutive in preorder and that have the same parent in M. The size of G is clearly upper bounded by the number of left extensions of maximal repeats of *T*. We use a bitvector `leafToMaxrep`[1..*|T |*] to mark such intervals of leaves in 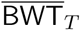, then we run-length compress it, i.e. we represent every maximal substring of the same character, as one occurrence of the character followed by the length of the substring (see e.g. [55] for more details). We also run-length encode the bitvector that represents open and closed parentheses in the topology of SLT_*T*_: a maximal run of open (or closed) parentheses corresponds to a unary path in SLT, i.e. to an edge between two maximal repeats. Finally, we run-length encode `mrSLT` and 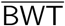: the latter can be shown to take a number of words that is upper bounded by the number of left extensions of maximal repeats [11]. Note that we don’t need bitvector `mrST`, since all nodes of *G* are maximal repeats except for the leaves.

We compute MS_*S,T*_ by scanning *S* from left to right: we use the run-length compressed 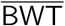 to decide whether a Weiner link is successful and to update the BWT interval of the current match. When a Weiner link by character c fails from the current interval in 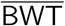, which corresponds to a node υ in 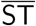, we know that the lowest ancestor of υ that has a Weiner link labelled by *c* must be a maximal repeat. Thus, we use bitvector `leafToMaxrep` to move in constant time to the interval of the lowest ancestor of υ that is a maximal repeat, and we try the Weiner link again from there. We can measure the string length of this ancestor, and of all its ancestors, since they are maximal repeats.

The probability of *S* according to the VOMM of *T* can be computed in a similar way. Indeed, we can use Lemma 1 to maintain, for each position *i* of *S*, the value MS[*i*] and the BWT interval of the matching statistics string *W* = *S*[*i -* MS[*i*] + 1..i]. Let υ be the locus of 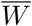 in 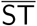. If contexts are maximal repeats, we use bitvector `leafToMaxrep` to move to the lowest ancestor of υ that is a maximal repeat (which might be υ itself). If contexts are left extensions of maximal repeats, we collapse leaves in *G* in a slightly different way: rather than collapsing into the same leaf all the leaves in a maximal preorder interval with the same lowest maximal repeat ancestor in 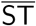, we collapse into the same leaf all leaves in the subtree of 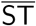 rooted at the same locus of a left-extension of a maximal repeat. This version of *G* still takes space proportional to the number of left-extensions of maximal repeats. We build `leafToMaxrep` correspondingly, and we build and run-length compress bitvectors `mrSLT` and context

These algorithms can benefit from few additional pruning strategies. For example, when computing MS with Lemma 1, one could further prune the topology of 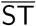 (but not of SLT) by keeping just the internal nodes of 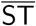 that have *a Weiner link that is not present in at least one of their children*, and by marking in a bitvector the corresponding nodes in SLT. This is because, when a Weiner link with character *S*[*i*+1] fails from the current matching statistics string *W*, the longest suffix of W that has a Weiner link labelled by *S*[*i* + 1] corresponds to a node of 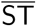 such that its child in the path to the locus of *W* does not have a Weiner link labelled by *S*[*i* + 1]. The internal nodes of 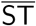 that are kept after this pruning are precisely the maximal repeats that are the infix of a minimal absent word of *T*, where a *minimal absent word* (MAW) of *T* is a string *W* that does not occur in *T*, but such that every proper substring of W occurs in *T* [20]. This pruning applies also to the scoring algorithm, if we just score character *S*[*i* + 1] with its emission probability from the matching statistics at position *i*, as done e.g. by [33]. Moreover, when contexts are left-extensions of maximal repeats, when the scoring function uses just the longest context that ends at the current position, and when the length of such context does not affect the score, it is easy to see that one can avoid storing the topology of SLT and the bitvectors to commute between 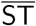 and SLT altogether.

A second way of reducing the size of our data structures consists in taking advantage of upper bounds on the length of contexts, which are frequently used in applications [18, 45, 46]. We focus here on using such constraint in matching statistics, leaving the immediate application to VOMM scores to the reader. Given an integer τ, we call *length-thresholded matching statistics* MS_*S,T,τ*_[1..*|S|*] an array such that MS_*S,T,τ*_ [*i*] equals MS_*S,T*_ [*i*] if MS_*S,T*_ [*i*] ≤τ, and it equals *–*1 otherwise.

### Lemma 2

*We can compute* MS*_S,T,τ_ using the setup of Lemma 1, but replacing the topologies of* 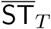 *and of* SLT_*T*_ *with subgraphs induced by maximal repeats of T of length at most* τ*, and by their left extensions.*

*Proof.* We proceed as in Lemma 1, but rather than considering the subgraph *G* of 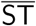 induced by *M* ∪ *L*, we consider the subgraph *G^′^* of 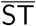 induced by *M*_τ_ ∪ *L*, where *M*_τ_ is the set of maximal repeats of length at most τ, and of all their children. We collapse onto the same leaf all the consecutive leaves of *G′* that have the same parent in *G′*. We also consider the subgraph of SLT induced by maximal repeats of length at most τ. We maintain the following invariant. At every position *i* of *S*, we know the value of MS_τ_ [*i*] and: (1) if MS_τ_ [*i*] ≠ –1, we know the interval of *W_i_* = *S*[*i -* MS[*i*] + 1..*i*] in 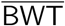; (2) if MS_τ_ [*i*]= *–*1, we know the interval in 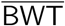 of a (not necessarily proper) suffix of *W_i_* of length greater than τ. We show that we can compute MS_τ_ [*i*] for every *i* while maintaining such invariant.

Assume that at some position *i* of *S* we are in case (2), and let υ be the (unknown) locus of *W_i_* in the full 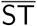. If υ can be extended to the right by *S*[*i*+ 1], then MS_τ_ [*i*+ 1] = *–*1, the right extension is possible also from the BWT interval of the suffix of *W_i_* that we have, and performing such extension maintains the invariant that we know the interval in 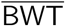 of a suffix of *W_i_*_+1_ of length bigger than τ. If υ cannot be extended to the right with character *S*[*i* + 1], but there is a Weiner link labelled by character *S*[*i* + 1] from the interval that we have, it means again that MS_τ_[*i*+ 1] = –1, and that υ is a descendant of the locus in 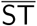 of our interval, thus taking the Weiner link preserves the invariant that we know the interval in 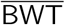 of a suffix of *W_i_*_+1_ of length bigger than τ. Finally, if υ cannot be extended to the right with character *S*[*i*+1], and if the interval we have cannot be extended to the right with *S*[*i* + 1] either, we use `leafToMaxrep` to convert the BWT interval that we have into an internal node υ′ of *G*′. Note that *|ℓ*(υ′)*|* ≥τ, but υ′ is not necessarily a maximal repeat. If υ′ does not have a Weiner link labeled by *S*[*i* + 1] either, then MS_τ_[*i*+1]≠–1, and we issue parent operations in the topology of *G′* that are identical to the parent operations we would have issued from v in the full 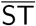, reaching eventually the same maximal repeat *w* that has a Weiner link labeled by *S*[*i* + 1] and whose length is at most τ. Otherwise, if υ′ has a Weiner link labeled by *S*[*i* + 1], then MS_τ_[*i* + 1] =–1, υ′ is necessarily a maximal repeat, and *w* is either υ′ itself or a descendant of υ′. Thus, by extending *υ′* (rather than *w*) to the right by *S*[*i*+ 1], we maintain the invariant that we know the interval in 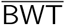 of a suffix of *W_i_*_+1_ of length greater than τ.

Finally, assume that at some position *i* of *S* we are in case (1). As long as *W_i_* can be extended to the right by *S*[*i* + *j*], we take a Weiner link and we update the value of MS_τ_ [*i* + *j*] for increasing *j*. Note that at each step we know the interval of MS[*i*+*j*], even when, for some *j*, a right extension leads to a string that is longer than τ. Finally, for some *j*, a right extension might not be possible. If MS[*i*+ *j*–1] > τ we do the same operations as in case (2), preserving the invariant. If MS [*i*+ *j –*1] ≤τ, the current interval contains a range of leaves of *G*′: we use `leafToMaxrep` to reach the lowest common ancestor of such leaves, which is necessarily a maximal repeat, and we continue from there.

Since frequency decreases monotonically with tree depth on a suffix tree, Lemma 2 can be adapted to compute the *frequency-thresholded matching statistics* MS_*S,T,τ*_[1..*|S|*], which is an array such that MS_*S,T,*_ [*i*] equals MS_*S,T*_ [*i*] if *f_T_* (*S*[*i -* MS_*S,T*_ [*i*] + 1..i]) ≥ τ, and it equals *–*1 otherwise.

## 4 Variants and extensions

We sketch the main ideas for supporting the variety of VOMMs and IMMs in Sections 2.2.1 and 2.2.2, omitting some technical details for brevity. We are not interested in achieving the fastest algorithm for each variant, but in showing that all variants can be implemented with just minor modifications to the non-pruned data structures of Section 2.3, and possibly to the pruned data structures of Section 3.

We already described in Section 3 how to support the scoring of [33]. To support the scoring scheme that uses the shortest right-deterministic suffix of *W_i_* as a context for position *i* [59, 19], it suffices to mark as contexts all nodes of 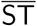 that are not maximal repeats, and such that their parent is a maximal repeat, and to proceed as in the case in which contexts are left-extensions of maximal repeats. Thus, this strategy can be implemented also with maximal repeat pruning. Recall that one of the statistical criteria used in Section 2.2.2 to mark a substring *W* as a context, checks whether the entropy of the emission probabilities from *W* is significantly larger than the entropy from the left-extensions of *W*. To support the symmetrical criterion of using the shortest *stable* suffix of *W_i_*, i.e. the shortest suffix that has *small* entropy compared to its left-extensions [61], it suffices to mark the loci of all stable substrings without a stable ancestor in 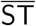, and to jump to one such locus from the locus of a matching statistics string *W_i_*. This strategy can thus be implemented with maximal repeat pruning as well. To enforce that no suffix of a context is itself a context [31, 54, 15], we can just unmark all nodes with a marked ancestor in 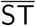. Enforcing that all prefixes of a context must be contexts (e.g. to make the Markov model memoryless: see e.g. [34]) does not guarantee that contexts are a subset of maximal repeats or of their left extensions. However, we can enforce that all prefixes of a context are suffixes of a context, by just adding leftmost maximal repeats or their left extensions to the set of contexts.

During construction we can compute the number of distinct characters in the BWT interval of every context, and we can store such numbers in an array of *O*(*m* log *σ*) bits, where *m* is the number of contexts. Given a query *S*, we can then compute the average number of right-extensions of all contexts that are used for scoring a position of *S* [37], by just accessing the array. The same approach can be used to access *σ*^ℓ^(*W*), *σ^r^*(*W*) or *σ^r^*(*W*, 1) for a context *W* at query time: together with *f*(*W*) and with the total frequency of single characters, such quantities are enough to implement a number of probability smoothings.

The recursive interpolation schemes of Section 2.2.1 can be implemented by iteratively taking parent operations in 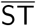, as described in Section 2.3. If the user provides an arbitrary weight function based on string length, we can implement the weighted mixing of suffix contexts by storing an array of prefix sums of log-weights: such array might not take too much space in practice if the string length of the longest edge of 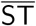 is small. If weights are constant, or if they are just a function of frequency and of the number of distinct characters to the right of a suffix of a context, they don’t change along an edge of 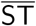. If they are a function of the length of a suffix of a context, and if such function is known in closed form [23], the contribution of all weighted probabilities along any edge of 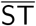 might even be computable in constant time. If we need the *maximum* emission probability among all suffixes of the longest context that are themselves contexts [37], we can just iterate calls to the lowest marked ancestor data structure. We can implement the IMM in Section 1 of the supplement in a similar way, by computing the values of *p*(*W*) during construction, and storing them in an array with one entry per left extension of a maximal repeat. We can then mark as context the locus υ in 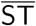 of all strings *aW* such that *f*(*aW*) is smaller than the given threshold and *W* is a maximal repeat. During scoring, we iteratively move from one context to the other using the lowest marked ancestor data structure.

To compute the backoff function *β*(*W, a*) of Section 2.2.1, we need to compute σ^ℓ^(*W* [2..*|W |*]*a*) for a context *W* [29]. Since contexts are maximal repeats, or left-extensions by one character of maximal repeats, we can decide whether 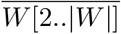 ends in the middle of an edge of 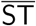 or not. If 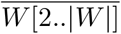 ends in the middle of an edge, then σ^ℓ^(*W* [2..*|W |*]*a*) is either one or zero, depending on whether a Weiner link with character a exists from the locus of 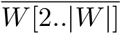 in 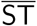. If 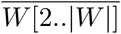 ends at a node of 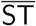, and if a Weiner link exists from such node with label *a*, then we need to count the number of distinct characters to the left of *W* [2..*|W |*]*a*, i.e. the number of children of the locus of 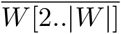 that have a Weiner link by character *a*. A similar method can be used to implement the Kneser-Ney smoothings in [52, 53, 51, 30]. Section 3 in the supplement describes how to compute some globals statistics on *T* using the same setup of data structures.

## 5 Experimental results

Our C++ implementation is based on the SDSL library [26] and is sequential: engineering it to use parallel threads falls outside the scope of this paper, and is discussed in Section 7 of the supplement. See Section 4 in the supplement for more implementation details.

We study the size and composition of our indexes in a number of datasets whose content and scale approximate current and possibly future applications of VOMMs. We focus on large datasets that were never tackled in previous works, and specifically: (1) The concatenation of all sequences in the latest reference assembly of the human genome in NCBI (5.9 billion characters). (2) The concatenation of all the approximately 8600 bacterial genomes currently in NCBI (34 billion characters). (3) The repetitive collections of bacterial genomes in the Pizza&Chili corpus^1^: these files range from approximately 100 to 400 million characters, and contain the concatenation of 23 genomes of *E. coli*, of approximately 35 genomes of *S. paradoxus* and of *S. cerevisiae*, and of approximately 78 thousand genomes of *H. influenzae*. (4) The concatenation of all non-redundant protein sequences in the NCBI RefSeq database ([41], 29 billion characters). (5) A repetitive dataset of proteins, built from the bacterial protein clusters in NCBI as described in Section 5 of the supplement (70 million characters). (6) The concatenation of all Illumina reads in a WGS metagenomic stool sample from the Human Microbiome Project ([36], 28 billion characters). (7) A prefix of the concatenation of all reads from a human individual in the Illumina Platinum project^2^ (33 billion characters). See Section 5 in the supplement for additional details on the datasets. The strings we use are long, but they are far from the longest ones that can be indexed by our implementation on a standard server with e.g. one terabyte of RAM (approximately 80 billion characters). For each string, we build: (1) the plain version of our index, with no pruning; (2) a version of the index in which topologies are pruned based on maximal repeats, and the BWT and the bitvectors are run-length encoded; (3) a version of (2), further pruned at depths that are powers of two. For concreteness, we select contexts using just the four-thresholds criterion of Section 2.2.2: see Section 5 in the supplement for details.

The size and composition of our indexes are summarized in Figure 3. The plain index takes approximately three bytes per character, with data structures for 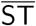 taking up the largest fraction of the total size, and bitvector context taking up approximately 16% of the total. Compression by maximal repeats saves between 30% and 50% of the size of the index in datasets that are not very repetitive, but it saves approximately 90% in repetitive collections like *S. cerevisiae*. In repetitive collections, bitvector `leafToMaxrep` takes approximately between 15% and 30% of the total size of the compressed index. Note that `leafToMaxrep` consists mostly of zeros, or mostly of ones, if there is little pruning or a lot of pruning, so it compresses well with run-length encoding. If there is a moderate amount of pruning, the distribution of zeros and ones is more even, and compression does not work as well. Further pruning by context length can shrink the compressed index to half of its size and beyond, but the the depth at which significant space reduction occurs depends on the dataset.

In large collections of short strings, like proteins or reads, or in datasets that are not very repetitive, maximal repeats are short and potentially few: rather than storing the topology of SLT and the bitvectors to move from 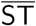 to SLT, it might be more space-efficient in practice to encode the length of every maximal repeat in pre-order. We experiment with the simplest possible scheme, i.e. storing each depth with a fixed number of bits that is just large enough to represent the largest length of a maximal repeat; we discuss more advanced methods in Section 8 of the supplement. Even with our simple encoding, explicit storage takes always less space than storing SLT, except in the human genome and in the concatenation of all bacterial genomes from NCBI (Figure 6 in the supplement). In the collection of *E. coli* genomes, explicit storage takes space comparable to the compressed SLT; otherwise, it allows one to save from 12% to 40% of the space of the compressed SLT.

Our VOMM implementation has marked advantages with respect to existing ones. As expected, the size of the index built by [5] depends on the number and length of contexts, so it produces smaller indexes than ours for restrictive settings of the thresholds. However, contrary to our data structure, index size blows up when thresholds are set to less restrictive values, becoming 10 times bigger than ours, or more (Figure 1): the limited scalability of pointer-based representations is the very motivation for research on space-efficient VOMMs. The code of [33], based on the suffix array, cannot index strings of length bigger than 80 million characters in practice, so we have to study its behavior on substrings of such length of all our datasets. Specifically, we take random sub-strings of the human genome, of the concatenation of all bacterial genomes, and of the concatenation of all non-redundant proteins, yielding non-repetitive strings; we use the full repetitive proteins dataset; and we take prefixes of the Pizza&Chili repetitive strings, which should still yield repetitive strings (albeit possibly less repetitive than the original). For non-repetitive strings, our index is between 3 and 4 times smaller than the competitor, and it becomes up to 11 times smaller after length pruning (Figure 2). In repetitive strings, our compressed index is from 10 to 60 times smaller than the competitor, and it becomes up to 500 times smaller after depth pruning (Figure 2). We remark again that such space savings should not be taken as upper bounds, since we are experimenting with short strings that might be less repetitive than the original datasets. We also note that, in addition to being larger, [33] supports just one type of score. The index in [53]^3^ takes from 3 to 3.3 bytes per character, regardless of the compressibility of the input. After removing precomputed counts used to speed up queries, the index takes between 2.6 and 3.1 bytes per character, which should approximate the size of the index in [52] and is comparable to our non-pruned data structure. Recall, however, that the competitor supports just an IMM with Kneser-Ney smoothing.

**Figure 1:**
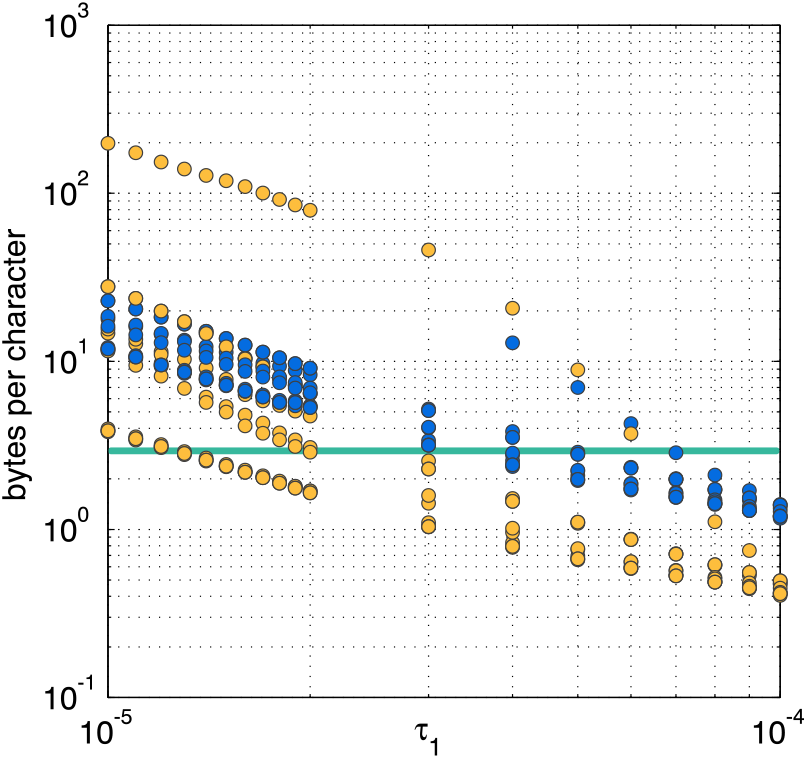
Size of trie representations of VOMMs as a function of τ_1_ (the minimum empirical probability of a context) in the four-thresholds context selection criterion of Section 2.2.2, over ten non-repetitive concatenations of proteins of length one million each. Decreasing τ_1_ implies increasing the number of contexts. Blue circles: the probabilistic suffix tree code in [5]; orange circles: the SPST code in [31]. Our implementation has a small variance, so it is represented just as a green line (averages) rather than as circles.

**Figure 2:**
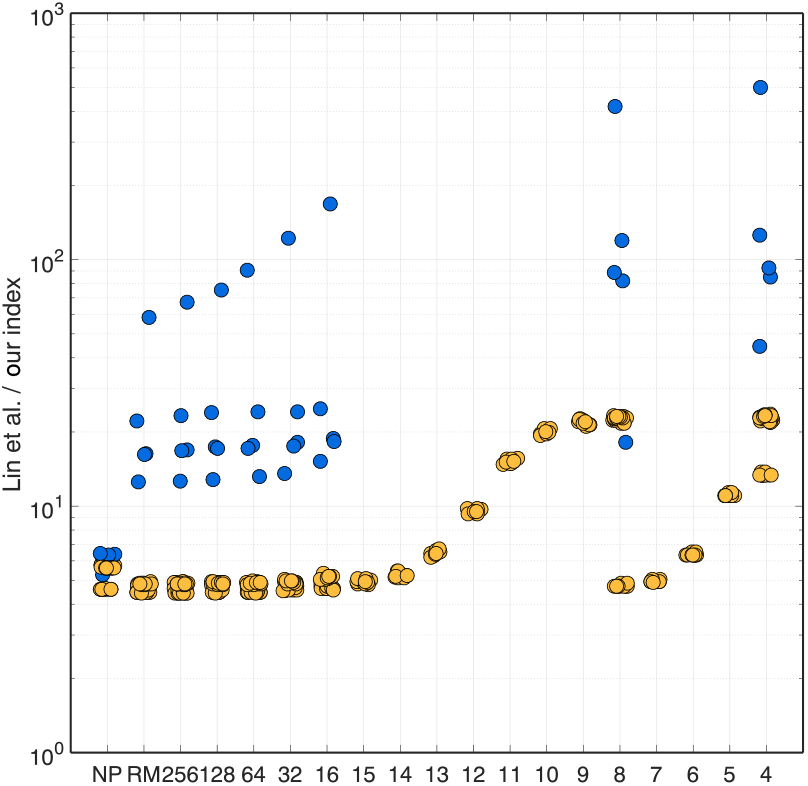
The size of our index compared to the size of [33], for strings of length at most 80 million. Orange circles: non-repetitive datasets; blue circles: repetitive datasets. See the text for more details. Note that the vertical axis has a logarithmic scale. Circles of the same color trace multiple curves because they belong to different datasets.

**Figure 3:**
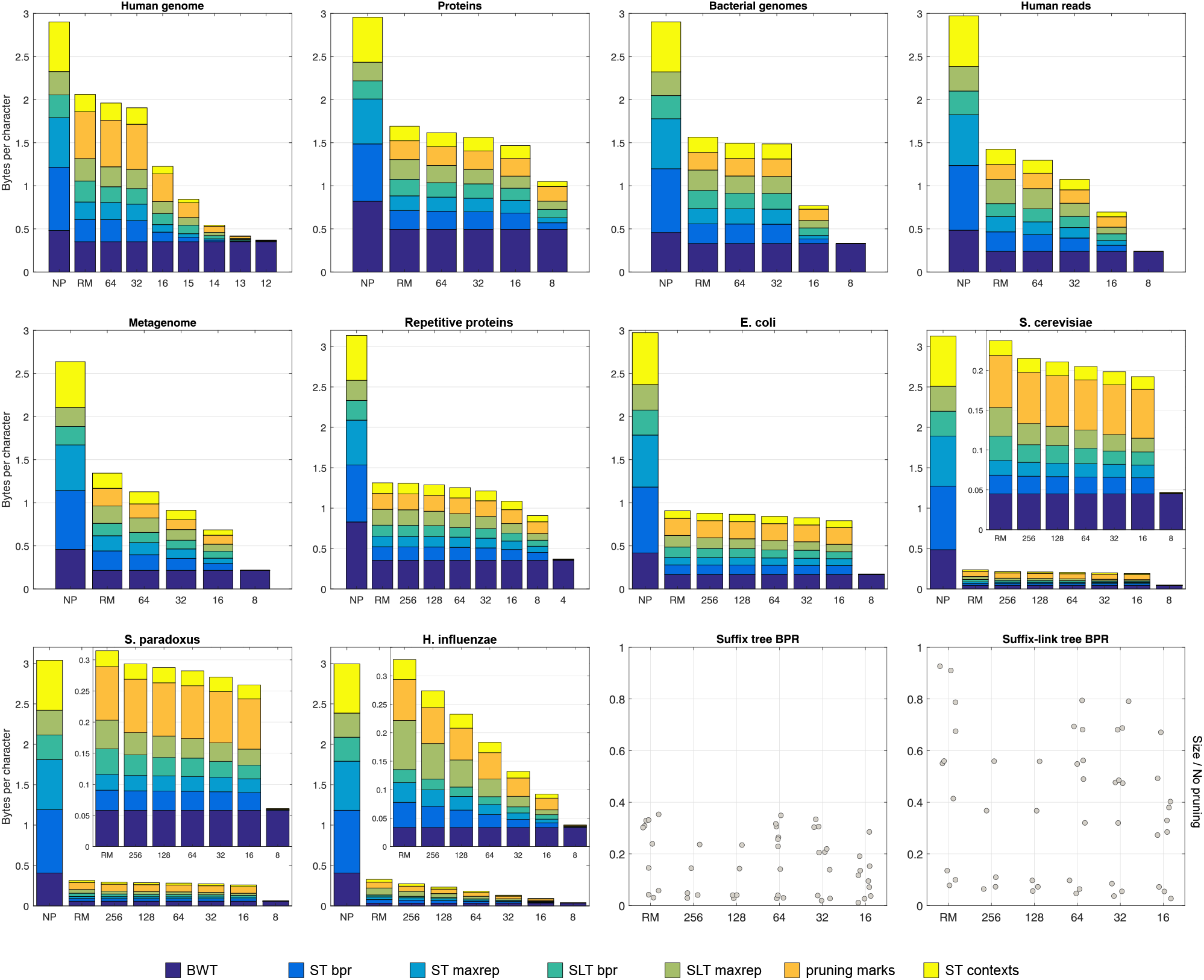
Size and composition of our indexes on large datasets. NP: no pruning; RM: run-length encoding (RLE) and maximal repeat pruning; numbers: RLE, maximal repeat pruning, and depth pruning. Pruning marks: all data structures associated with pruning 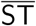 and SLT, including `leafToMaxrep`. The two dot plots on the bottom-right panels show the size of the balanced parentheses representations of 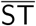 and SLT, divided by their size without pruning (bitvectors context, `mrSLT` and `mrST` have similar plots).

Optimizing index construction is outside the scope of this paper. However, our sequential construction algorithms are reasonably fast and space-efficient in practice: see Section 6 in the supplement for details and for a comparison to the competitors. Maximizing the speed of scoring functions is also outside the scope of this paper. Studying scoring time is interesting, however, since we expect to be paying our space savings in this currency. We report a detailed analysis of scoring time in Section 7 of the supplement. Here we just mention that non-recursive scoring on our index with maximal repeats pruning is indeed approximately 5-6 times slower than the same scoring with our non-pruned index; however, pruning the compressed index further by maximum length of a context makes scoring up to 2 times *faster*. Our non-pruned index is between 60 and 35 times slower than the trie-based index in [5], and it is between 3 and 12 times slower than [33] to compute the specific score of the competitor, depending on the dataset.

## Acknowledgements

We thank V. Mäkinen for guidance, E. Shareghi for help with the code of [53], and O. Gonzalez and P. Steinbach for help with the MPI-CBG cluster.

## 1 IMMs and maximal repeats

We show that the IMM in [29] performs a linear combination of the emission probabilities of just *the left extensions by one character of maximal repeats*. Recall from Section 2.2.1 in the paper that such IMM assigns the following score to observing character *b* after context *W*: *P*〰(*b|W*) = λ(*W*) *P*(*b|W*) + (1 *–* λ (*W*)) *P*〰(*b|W* [*2*..*|W |*]) if *|W |* > 0, and *f*(*b*)/*|T |* otherwise. If *f*(*W*) is at least a given threshold, then,λ(*W*) is set to one and recursion stops. Otherwise, *λ*(*W*) is set to either zero or, λ′(*W*) = c *·* (1 *– p*(*W*)) *f*(*W b*), where *c* is a constant and *p*(*W*) is the fraction of mass of the x distribution to the right of the Pearson’s chi-squared test statistic *x*(*W*). Such statistic uses the frequencies of *W_a_* as observed frequencies, and the frequencies of *W* [2..*|W |*]*a* as expected frequencies, for every *a* ∈ Σ, i.e. *x*(*W*)=*f*(*W*). ∑_*a*_∈ Σ^*r*^_(W[2..|W|])_(*P*(*a*|*W*)–*P*(*a*|*W*[2..|*W*|]))^2^/*P*(*a|W*[*2..*|*W*|]). The IMM chooses between setting, λ(*W*) to zero or to λ′(*W*) by comparing 1– *p*(*W*) to a positive, user-defined threshold. Note that, if *W* [2..*|W |*] if not left-maximal, *P*(*a|W*) coincides with *P*(*a|W* [2..*|W |*]) for any *a*, thus *x*(*W*) = 0. Similarly, if *W* [2..*|W |*] is always followed by character *b*, the summation in *x*(*W*) runs just over *b*, and both *P*(*b|W*) and *P*(*b|W* [2..*|W |*]) are equal to one, thus again *x*(*W*) = 0. When *x*(*W*) = 0, we also have that *p*(*W*) = 1, 1 –*p*(*W*) = 0, and λ (*W*) = 0.

## 2 Contexts as left extensions of maximal repeats

We describe how to modify the algorithm in Section 2.3 of the paper to handle contexts that are left extensions by one character of maximal repeats. Assume again that we mark the locus of all such contexts in the topology of 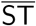. After taking a sequence of successful Weiner links from the first position of *S*, assume that we are currently at position *i* of *S*. We go to the locus υ of the reverse of *W* = *S*[1..*i*] in 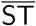 and, if υ is marked, we use the emission probability from υ to score *S*[*i* + 1]: this is equivalent to using the emission probability of the context *S*[*i*–*ℓ*(υ′)..i], where υ′ is the parent of υ in 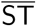. If υ is not marked, we move to its lowest marked ancestor υ′. When we back off from a node *w* to a node *w*′ of 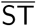, where *w* does not emit *S*[*i* + 1] and *w*′ if the lowest marked ancestor of *w* that emits *S*[*i* + 1], we adapt the algorithm to take into account that now we are considering contexts *S*[*i –ℓ*(*u*)..*i*],…, *S*[*i –ℓ*(*u^′^*)..*i*], where *u* is the parent of *w* and *u^′^* is the parent of *w*′.

## 3 Statistics on the training data

Note that, once we have built our data structures on *T*, we can sample a string *S* from the VOMM using the scoring algorithm in Section 2.3 of the paper, with just the topology of 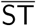, the BWT of the reverse, and the lowest marked ancestor data structure: indeed, we can start from a random context, and iteratively sample a Weiner link at random, based on the emission probabilities of the current context, and then move from the locus of the resulting extension to its lowest ancestor in 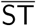 that is a context.

The same setup can be used to compute a number of global statistics on *T*, for example a maximum-likelihood estimate of the order of the fixed-order Markov chain that generated *T*: this is the value of *k* that minimizes –log *P*̂_k_(*T*) + (σ–1)σ^*k*^ log (*|T |– k* + 1)/2, where *P*̂_k_(*T*) = ∏_*W* ∈Σ_^*k*^ *f*(*W a*)>0(*f*(*W a*)/*f*(*W*))^*f*(*Wa*)^ [14]. To compute *P*̂_k_(T) for a specific value of *k*, it suffices to mark as contexts just the loci of *k*-mers in 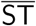, and to score *T* against its VOMM. The same holds for computing the *k-th order Markov predictability* of *T*, defined as (1/(*|T |–k*)) *∑*_W ∈Σ_^*k*^, *∑_a_* _∈Σ\{γ(W)}_ *f*(*Wa*)where γ (*W*) = argmax{a ∈ *f*(*Wa*)> 0}, for a user-specified range of values of *k* [15]. This is the fraction of errors while predicting the characters of *T* using contexts of fixed length *k*, predicting as the next character, at each position of *T*, the character with largest relative frequency with respect to the current *k*-mer.

## 4 Details on the implementation

We build the BWT with the divsufsort library^1^. We use the rank data structure `rank_support_v` and the code for select operations on bitvectors `select_support_mcl` from the SDSL library [18], and we represent the topology of 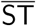 and of SLT explicitly with balanced parentheses, using `bp_support_g`. We build most data structures using our own implementation of the bidirectional BWT index^2^, and we use portions of the `lz-rlbwt` code^3^ from [10] for the run-length-encoded BWT. One could plug in other implementations to build the indexes, for example [2] for the suffix tree topology; we haven’t experimented with such alternatives since we do not try to optimize construction. The implementation ships with comprehensive unit tests on randomized strings to ensure correctness.

When comparing our implementation to the code in [4], we turn off computing and storing statistics in the competitor’s source, and we compile it with `gcc-O3`.

## 5 Details on the datasets

Our metagenomic stool sample is file SRS301868 at https://portal.hmpdacc.org/files/91319e642fdd8a6e3b059cfb058cc4aa, and our human reads from the Illumina Platinum project are from run `ERR194146`, file `ERR194146_1.fastq.gz`, from https://www.ebi.ac.uk/ena/data/view/PRJEB3381

When building each dataset, we concatenate its sequences using a separator that is not in the alphabet, replacing runs of undetermined characters with a single occurrence of the separator. We build the dataset of repetitive proteins as follows. We sort all bacterial protein clusters in NCBI^4^ (which are sets of protein sequences with high pairwise alignment scores) by decreasing number of sequences in a cluster. For every such cluster, we concatenate all sequences in it, in order, until we reach approximately 70 million characters.

In all datasets we select contexts using the four-thresholds criterion, with the parameters τ_1_ = 10^*-*4^, τ_2_ = 10^*-*3^, τ_3_ = 0.952, τ_4_ = 1.05 used for proteins in [5] (similar values are also used in [23, 3, 22, 6]). Using different criteria does not fundamentally change our results.

## 6 Index construction

We sketch some techniques that we observed are effective for building VOMMs in practice.

### 6.1 Balanced parentheses

We build the balanced parentheses representation of the full topology of 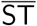 (without pruning) by iterating over all left-maximal substrings of *T* with 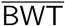, as described in [7, Section 4.1]. Specifically, we compute the lexicographic range [*i_W_*..*j_W_*] in 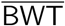 of every left-maximal substring W. Let open[1..*|T|*] (respectively, close[1..*|T|*]) be an array that stores the number of substrings W such that *i_W_* = *i* (respectively, *j_W_* = *j*). Given such arrays, one could build the balanced parentheses representation of 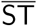 by iteratively printing open[*i*] open parentheses followed by close[*i*] closed parentheses, for all values of *i*. To build open and close we could just increment open[*i_W_*] and close[*j_W_*] for each traversed *W*. Such vectors take *|T|* log *|T|* bits of space, so it is desirable to make them smaller. The first idea to reduce their space in practice is based on the fact that min(open[*i*], close[*i*]) < 2 for every *i*, since otherwise 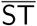 would have a unary path. We can thus encode both open[*i*] and close[*i*] in a single array of counters openClose[1..*|T|*], using e.g. the following encoding: a zero at position i means that there is no open and no closed parenthesis at *i*; a one means that there is just one open parenthesis; a two that there is just one closed parenthesis; a three that there are one open and one closed parenthesis; an even number *n* ≥ 4 means that there are *n*/2 open parentheses and one closed parenthesis; an odd number *n* ≥ 5 means that there are (*n* – 1)/2 closed parentheses and one open parenthesis. Note also that in practice most counters are small, thus we use a two-level scheme, in which an array stores values of length at most b bits, and a hash table stores all other values^5^. We can store approximately 2^*b-*1^ parentheses per position of the array: if we need more, we mark the position as *saturated* and we add a new counter to the hash table, where keys are BWT positions and values are 64-bit integers. We could use a similar algorithm to build the balanced parentheses representation of the full topology of SLT (without pruning), except that now we would iterate over all right-maximal substrings of *T* using BWT, incrementing counts in 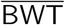 using the bidirectional BWT index to synchronize the two intervals. However, since the stack of the iterator is small in practice, we just perform a preorder traversal of SLT and append open and closed parentheses, building bitvector `mrSLT` at the same time.

To build bitvector `mrST`, we traverse the topology of 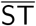 in any order, we use the topology to get the interval [*i*..*j*] of each node in 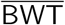, and we check the number of ones in the interval [*i* + 1..*j*] inside an additional bitvector `diff`[1..*|T|*] such that `diff`[*i*] = 1 `iff` 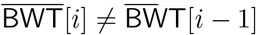 [11].

We build the data structures in Lemma 1 of the paper as follows. We build the balanced parentheses representation of the topologies of 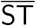 and SLT like in the non-pruned case by traversing SLT using the birectional BWT index, but keeping only nodes that are maximal repeats or left-extensions of maximal repeats. We also mark the required bits in `leafToMaxrep` during the traversal.

### 6.2 Complexity and comparison to the competitors

We can measure most quantities used in Section 2.3 of the paper to decide if a node of 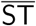 is a context, within the same time and space budget as traversing the tree using 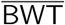. Thus, for most variants, construction takes *O*(*|T|* log *σ*) time. In general, construction takes *O*(*υ|T|* log *σ*) time, where v is the time to decide whether a node of 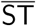 is associated with a context.

Computing the contexts for a different setting of the context selection thresholds amounts just to a traversal of 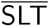, thus *re-training a VOMM that was already built is significantly faster than training one from scratch*. For the same reason, *storing multiple VOMMs trained on the same dataset with different settings takes significantly less space than the sum of the individual VOMMs*.

On an Intel Xeon E7-4830v3 at 2.10 GHz with one terabyte of RAM, our sequential construction algorithm takes approximately ten hours to index the human genome, and between three and four days to index all bacteria, all proteins, the metagenome, and the read set. Trie-based data structures are from 5 to more than 290 times slower to build than our index in practice, depending on the number of contexts (Figure 1). Building the index of [24] is approximately 3 times faster than building our non-pruned index, and 5 times faster than building our pruned index, except for aggressive depth pruning (Figure 2), but our construction algorithm uses approximately half the space of the competitor. Once our index is built, creating one with a different context selection criterion is faster than building *the competitor* from scratch, and it takes half the space required for building our index from scratch. Building the index in [32] takes between 5 and 9 bytes per character, which is comparable to our construction, and between 1.1 and 3.2 microseconds per character, which is faster than or comparable to our non-pruned index.

Note that, once our indexes are built, the user can also discard the training data and use the index itself to reconstruct *T* if needed. Note also that all construction algorithms could be parallelized, by performing parallel traversals of 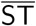 and of SLT (see e.g. [1]).

## 7 Scoring time

We implement just basic versions of each scoring criterion, paying attention only at minimizing the number of conversions between node identifier in a topology and corresponding BWT interval. Clearly the query S does not need to be kept in memory but can be streamed from disk.

### 7.1 Comparison to the competitors

As mentioned, our non-pruned index is between 60 and 35 times slower than the trie-based index in [4] (Figure 3). Contrary to the competitors, however, scoring time *decreases* when the number of contexts increases: this is likely due to the fact that, when only few contexts are selected, most matching statistics strings need to issue a lowest marked ancestor query to jump to their context, whereas when many contexts are selected, it is more likely for a matching statistics string to already be a context, and thus to avoid a lowest marked ancestor query.

Recall that the implementation in [24] cannot index strings longer than approximately 80 million characters. Thus, we compare the speed of scoring with our index and with the competitor, by using the following small datasets as indexes: (1) a random subset of all NCBI proteins, of length 80 million; (2) the set of repetitive proteins described above; (3) a random subset of bacterial genomes, of length 80 million; (4) a prefix of the *H. influenzae* file, of length 80 million. We use such datasets to cover the cases of repetitive and non-repetitive indexes. We use as queries ten random subsets of all NCBI proteins, and ten random subsets of all bacterial genomes, of length 500 million each. Recall also that the competitor does not allow one to use contexts to score a query; instead, it just computes the ratio between the frequency of *S*[*i-* MS[*i*]+ 1..*i*] and the frequency of *S*[*i –* MS[*i*]+ 1..*i –* 1] for each *i*. We implement such criterion using an even smaller data structure, since we don’t need to measure string depths. Results are in Figure 5. We don’t compare scoring time to [32], since the latter supports just one scoring function which is significantly different from the ones we consider.

### 7.2 Recursive scoring

It is interesting to compare recursive and non-recursive scoring time when the number of contexts increases. As before, we decrease threshold *τ*_1_ in the four-thresholds selection criterion of Section 2.2.2 in the paper. When *τ*_1_ is large, recursive and non-recursive scores take similar time, since we are likely selecting very short contexts, which are probably followed by most characters of the alphabet, and thus recursion stops soon (see Figure 5). As expected, when *τ*_1_ decreases, the recursive score becomes slower, since more substrings are marked as contexts, thus the matching statistics string at each position of the query tends to match a longer context, which is more likely to emit just few characters, and thus recursion is more likely to continue upwards in 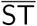. We observe the recursive score being up to 3.5 times slower than the non-recursive score, and the gap widens as τ_1_ decreases.

### 7.3 Speeding up scoring

Slower query times could be mitigated in practice by applying optimizations from matching statistics (like those in [9], some of which are implicitly enabled by our pruned topologies already), by precomputing counts that are too expensive to evaluate at query time (as done e.g. in [21, 30, 31, 32]), and by taking advantage of the large number of cores that are standard in current servers. Scoring is indeed embarrassingly parallel in most applications, where the dataset to be queried is a large number of short strings, like sequencing reads or proteins. If *S* is one long string, one can still split it in uniform blocks and parallelize matching statistics.

## 8 Explicit storage of string depths

Figure 6 shows the advantages of a naive explicit encoding of the lengths of maximal repeats. Alternatively, we could use a multilevel scheme like *directly addressable codes* [12], which have already been used e.g. by [32] for storing precomputed counts. Specifically, a first array could contain one byte for every maximal repeat in preorder, and the first bit in such byte could mark whether the length of the maximal repeat is longer than 2^7^. A second array stores again one byte for every maximal repeat of length greater than 2^7^, and uses again the first bit to mark whether the length is greater than 2^14^. A third array contains the remaining log *d* –14 bits that encode the length of every maximal repeat of length greater than 2^14^, where *d* is the maximum length of a maximal repeat. To move from one array to the other, we could store partial rank information every few bytes.

Another alternative could be to simply store the difference between the string depths of maximal repeats that are consecutive in preorder, using two bitvectors add and subtract, each containing exactly a number of ones equal to the number of maximal repeats: if the difference between the string depths of node *i* and *i–*1 in preorder is equal to *d*, we append *d* zeros followed by a one to add, and if it is equal to *–d* we append *d* zeros followed by a one to subtract.

Note that storing depths explicitly might reduce query time, since fewer operations are needed to access the length of a maximal repeat. In practice we observe negligible speedups in all datasets, even with recursive scores, suggesting that string depth retrieval is not a bottleneck of score computation.

## 9 Background on string indexes

We briefly summarize here some background notions on data structures that are required for understanding the paper.

Let *T* ∈ [1..*σ*]^*n-*1^# be a string. A *rank query* rank(*T*, *a*, *i*) returns the number of occurrences of character a in *T* up to position *i*, inclusive. A *select query* select(*T*, *a*, *i*) returns the position of the *i*-th occurrence of *a* in *T*. See e.g. [17]. The *suffix array* SA_*T*_ [1..*|T|*] of *T* is the vector of indices such that *T* [SA_*T*_[*i*]..*|T|*] is the *i*-th smallest suffix of *T* in lexicographic order. The Burrows-Wheeler transform of *T* is the string BWT_*T*_ [1..*|T|*] satisfying BWT_*T*_ [*i*] = *T* [SA_*T*_ [*i*] *-* 1] if SA_*T*_ [*i*] > 1, and BWT_*T*_ [*i*] = # otherwise [13]. While SA_*T*_ takes *|T|* log *|T|* bits, BWT_*T*_ takes *|T|* log *σ* bits, i.e. the same number of bits needed to store *T*. A substring *W* of *T* can be represented as a lexicographic interval [*i*..*j*] of suffixes in both SA_*T*_ and BWT_*T*_. Array *C*[0..*σ*] stores in *C*[*a*] the number of occurrences in *T* of all characters strictly smaller than *a*, i.e. the sum of the frequency of all characters in set {#, 1, …, *a -* 1}. Clearly *C*[0] = 0, and *C*[*a*] + 1 is the position in SA_*T*_ of the first suffix of *T* that starts with character *a*. The *wavelet tree* is a data structure that represents a string *T* in *|T|* log *σ*(1 + *o*(1)) bits and supports rank, select, and access operations on its characters in *O*(log *σ*) time (see e.g. [19]). The combination of BWT with rank support and *C* array is known as *FM-index*, and it enables a *backward step*, i.e. moving from the interval [*i*..*j*] of a substring *W* of *T*, to the interval [*i′*..*j′*] of its left extension *aW*, where a is a character [16].

We also require familiarity with the notion and usages of the *suffix tree* ST_*T*_ = (*V*, *E*) of *T* # [33]: see e.g. [20] for an overview. We denote by 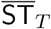 the suffix tree of 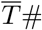, and we denote by *ℓ*(*υ*) the string label of a node *υ* ∈ *V*, i.e. the string obtained by concatenating the labels of all edges in the path from the root of the tree to *υ*. Here we just recall that a substring *W* of *T* is right-maximal (respectively, left-maximal) iff *W* = *ℓ*(*υ*) for some internal node *υ* of ST_*T*_ (respectively, for some internal node *υ* of 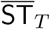), and that a node *υ* ∈ *V* with *ℓ*(*υ*) = *aW* for some character *a* ∈ [0..*σ*] points to a node *w* ∈ *V* with *ℓ*(*w*) = *W* by a *suffix link* labeled by a. Suffix links and internal nodes of ST_*T*_ form a trie, called the *suffix-link tree* of *T* and denoted by SLT_*T*_. Inverting the direction of all suffix links yields the so-called *explicit Weiner links*. Given an internal node *υ* and a symbol a ∈ [0..*σ*], it might happen that string *aℓ*(*υ*) occurs in *T*, but is not right-maximal, i.e. it is not the label of any internal node of ST: all such left extensions of internal nodes that end in the middle of an edge are called *implicit Weiner links*. An internal node *υ* of ST can have more than one outgoing Weiner link, and all such Weiner links have distinct labels: in this case, *ℓ*(*υ*) is a maximal repeat. We call 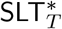 a version of SLT_*T*_ augmented with implicit Weiner links and with nodes corresponding to their destinations. A maximal repeat *W* of *T* is called *leftmost* if it is not the proper suffix of any other maximal repeat of *T*. Since taking the suffix of a string preserves right-maximality, the set of all maximal repeats coincides with the set of all ancestors in 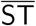 of leftmost maximal repeats. It is easy to see that there is a bijection between the set of branching nodes of SLT* and the nodes of 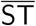 that correspond to maximal repeats, in which the leaves of SLT are mapped to the nodes of 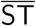 that correspond to leftmost maximal repeats. The suffix-link tree (a *trie*) is thus a *subdivision* of the subgraph of 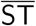 induced by maximal repeats (a *compact tree*). See Section 2.1 in [8] for a more thorough explanation.

The topology of an ordered tree with *n* nodes can be represented using 2*n* + *o*(*n*) bits, as a sequence of 2*n* balanced parentheses built by opening a parenthesis, by recurring on every child of the current node in order, and by closing a parenthesis [25]. Let *id*(*υ*) be the rank of a node v in the preorder traversal of the tree. Given the balanced parentheses representation of the tree encoded in 2*n* + *o*(*n*) bits, one can build a data structure that takes 2*n*+*o*(*n*) bits, and that supports the following operations in constant time [26, 28, 27]:

- `child`(id(*υ*), *i*): returns id(*w*), where *w* is the *i*th child of node *υ* (i ≥ 1), or ∅ if *υ* has less than i children;
- `parent`(id(*υ*)): returns id(*u*), where *u* is the parent of *υ*, or ∅ if *υ* is the root of *T*;
- `lca`(id(*υ*), id(*w*)): returns id(*u*), where *u* is the lowest common ancestor of nodes *υ* and *w*;
- `leftmostLeaf`(id(*υ*)), `rightmostLeaf`(id(*υ*)): returns one plus the number of leaves that, in the preorder traversal of *T*, are visited before the first (respectively, the last) leaf that belongs to the subtree of *T* rooted at *υ*;
- `selectLeaf`(*i*): returns id(*υ*), where *υ* is the *i*-th leaf visited in the preorder traversal of *T*;
- `depth`(id(*υ*)), `height`(id(*υ*)): returns the distance of *υ* from the root or from its deepest descendant, respectively.

This data structure can be built in *O*(*n*) time and in *O*(*n*) bits of working space.

Note that such operations allow converting in constant time between an interval in BWT_*T*_ and the corresponding node identifier in the balanced parenthesis representation of ST_*T*_ (by selecting the corresponding leftmost and rightmost leaves and taking their lowest common ancestor), and vice versa (by counting the number of leaves under a node).

---

**Figure 1:**
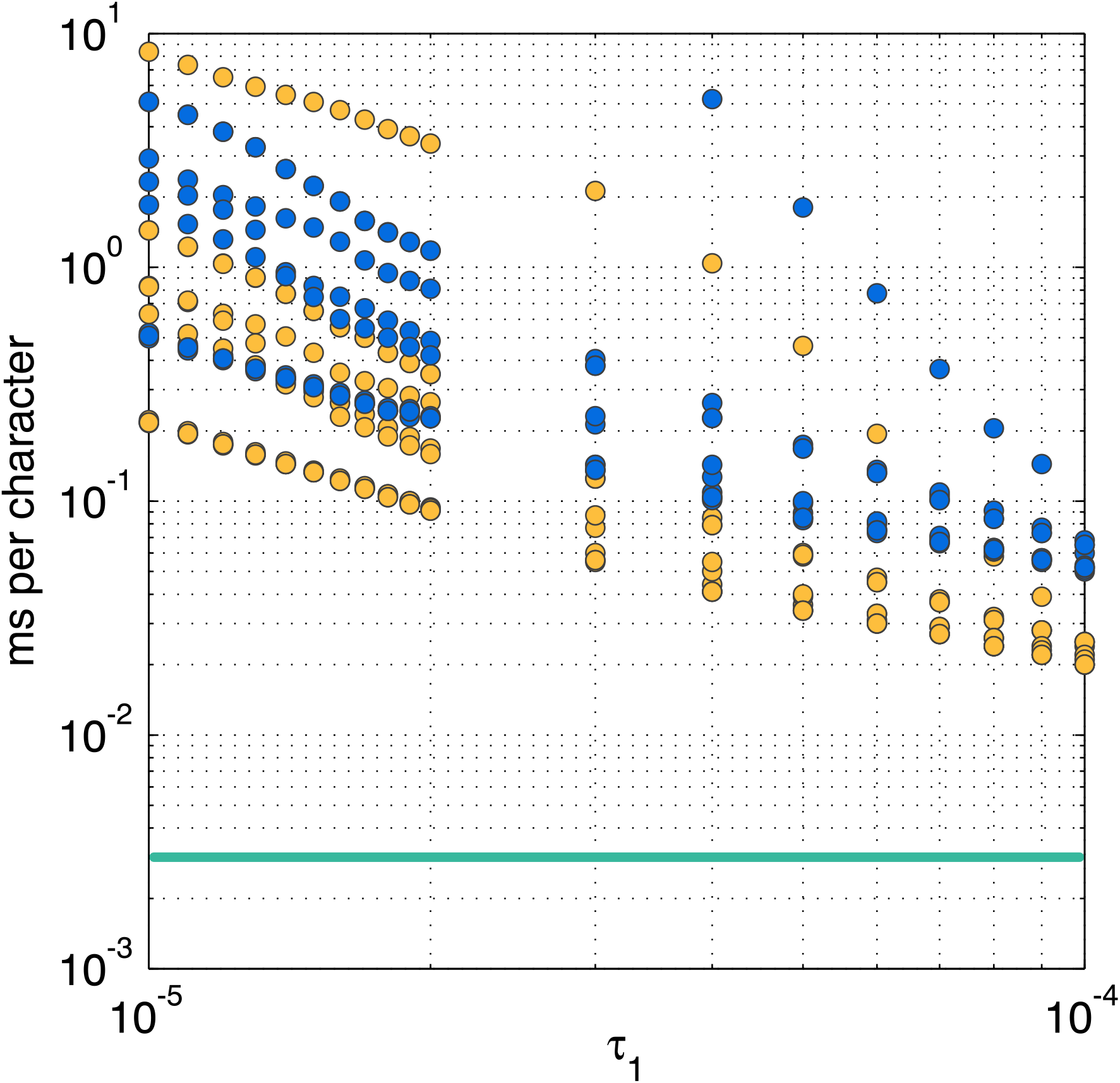
Size of trie representations of VOMMs as a function of τ_1_ (the minimum empirical probability of a context) in the four-thresholds context selection criterion of Section 2.2.2, over ten non-repetitive concatenations of proteins of length one million each. Decreasing τ_1_ implies increasing the number of contexts. Blue circles: the probabilistic suffix tree code in [5]; orange circles: the SPST code in [31]. Our implementation has a small variance, so it is represented just as a green line (averages) rather than as circles.

**Figure 2:**
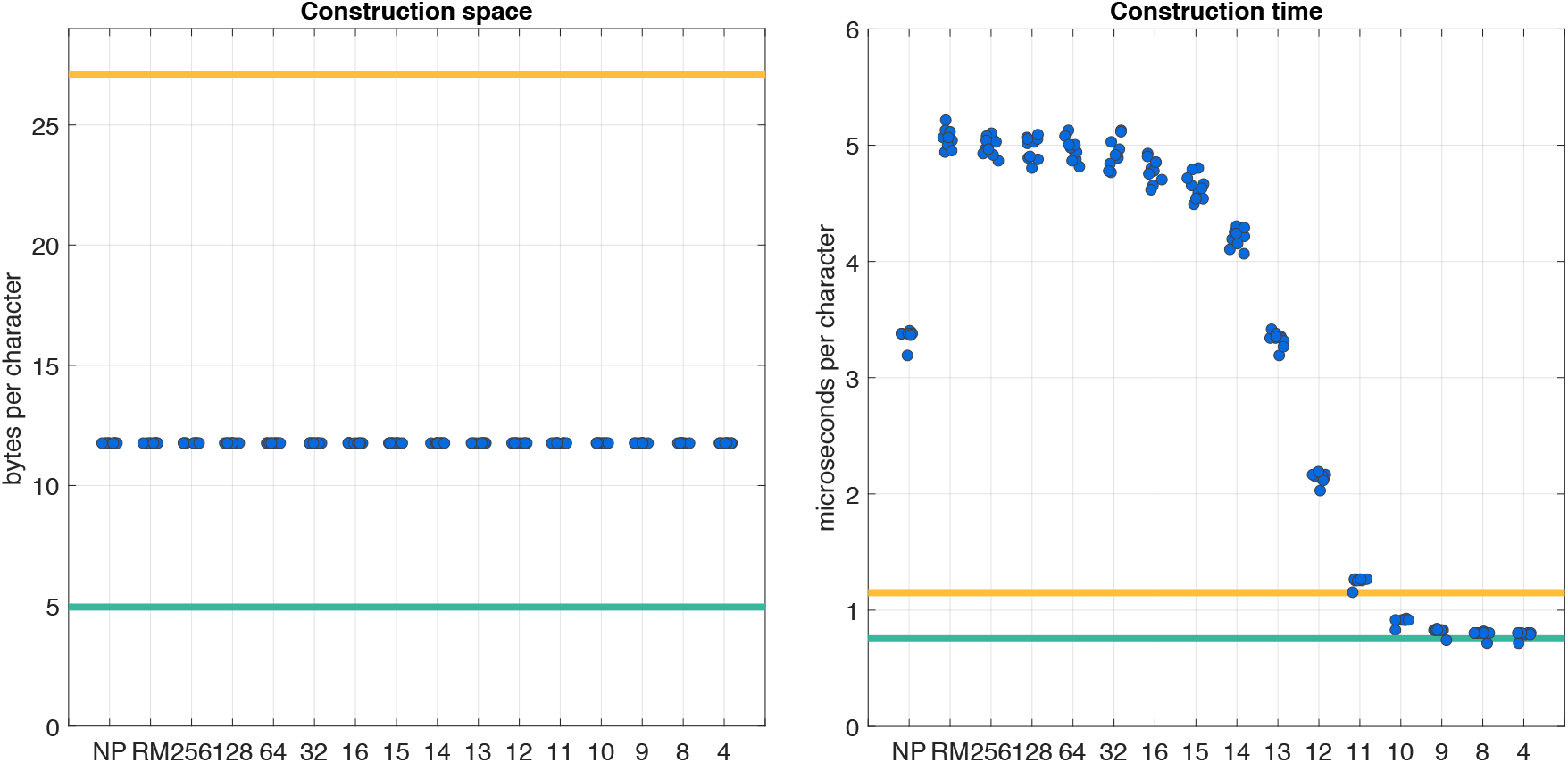
The size of our index compared to the size of [33], for strings of length at most 80 million. Orange circles: non-repetitive datasets; blue circles: repetitive datasets. See the text for more details. Note that the vertical axis has a logarithmic scale. Circles of the same color trace multiple curves because they belong to different datasets.

**Figure 3:**
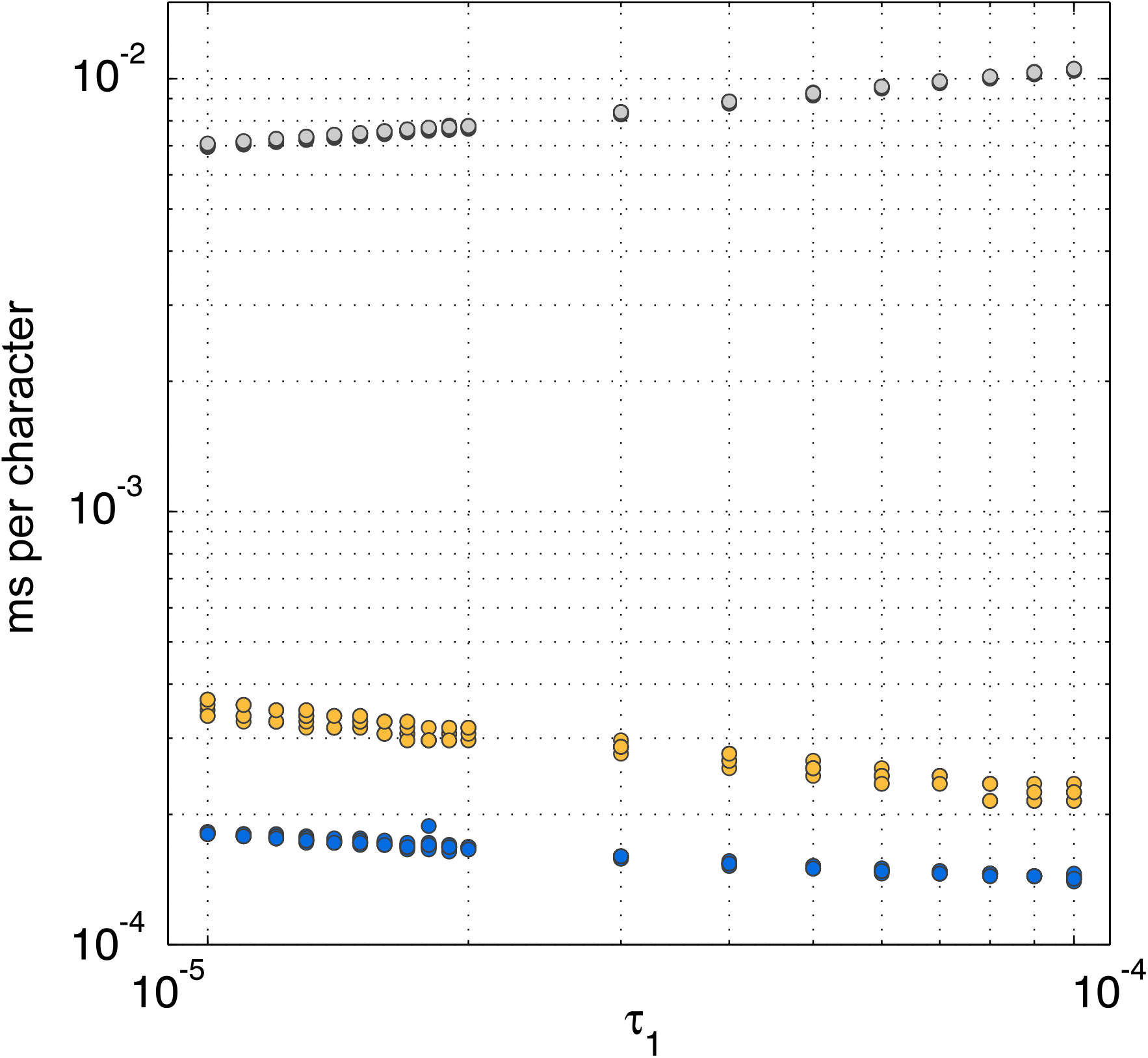
Size and composition of our indexes on large datasets. NP: no pruning; RM: run-length encoding (RLE) and maximal repeat pruning; numbers: RLE, maximal repeat pruning, and depth pruning. Pruning marks: all data structures associated with pruning 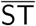 and SLT, including `leafToMaxrep`. The two dot plots on the bottom-right panels show the size of the balanced parentheses representations of 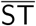 and SLT, divided by their size without pruning (bitvectors context, `mrSLT` and `mrST` have similar plots).

**Figure 4:**
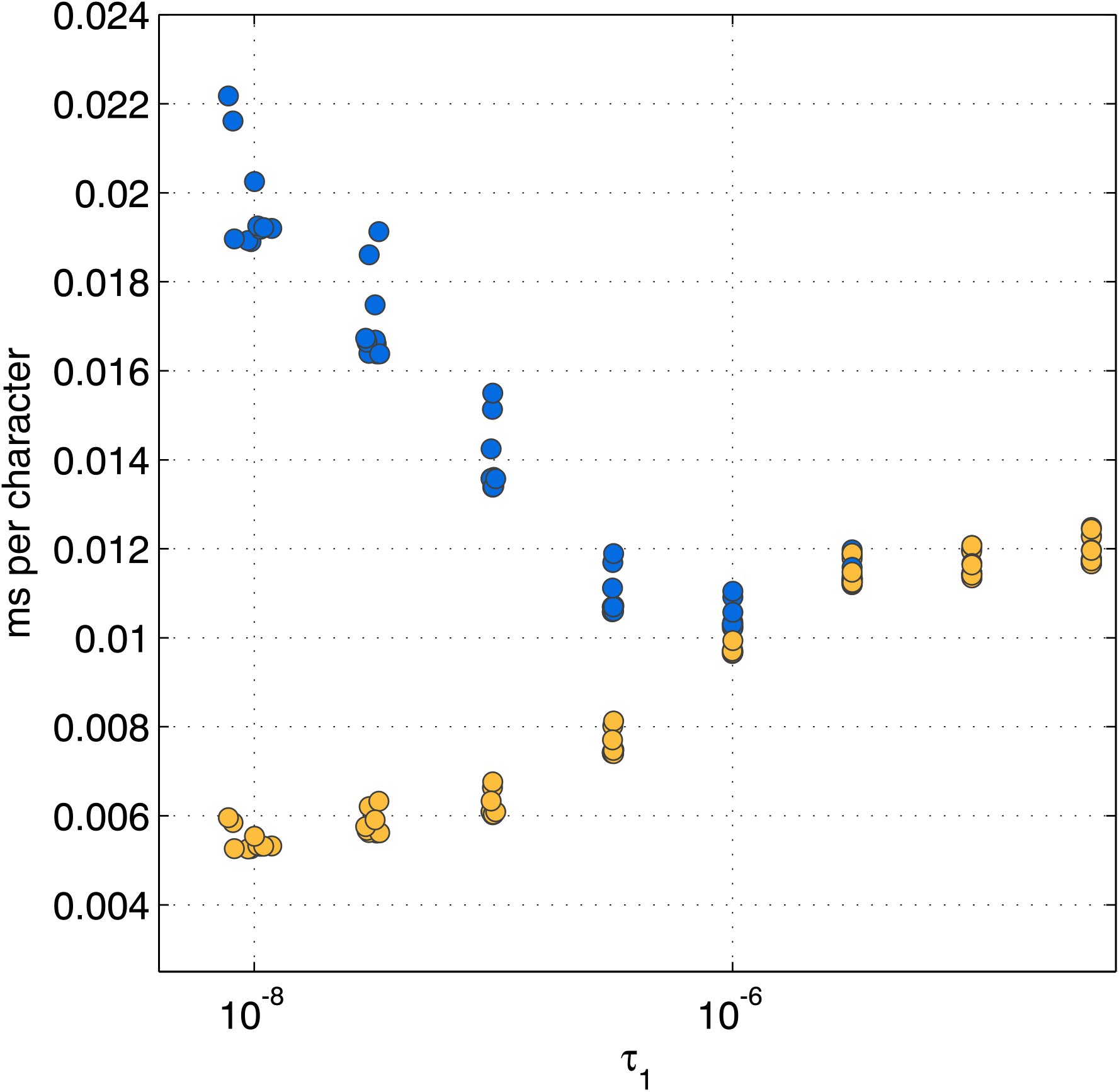
Recursive (blue circles) and non-recursive (orange circles) scoring in our implementation, as a function of *τ*_1_ in Section 2.2.2. of the paper (horizontal axis, logarithmic scale). Queries: ten non-repetitive concatenations of proteins of length 500 million. Index: one non-repetitive concatenation of proteins of length 80 million. Experiments with repetitive proteins yield similar results.

**Figure 5:**
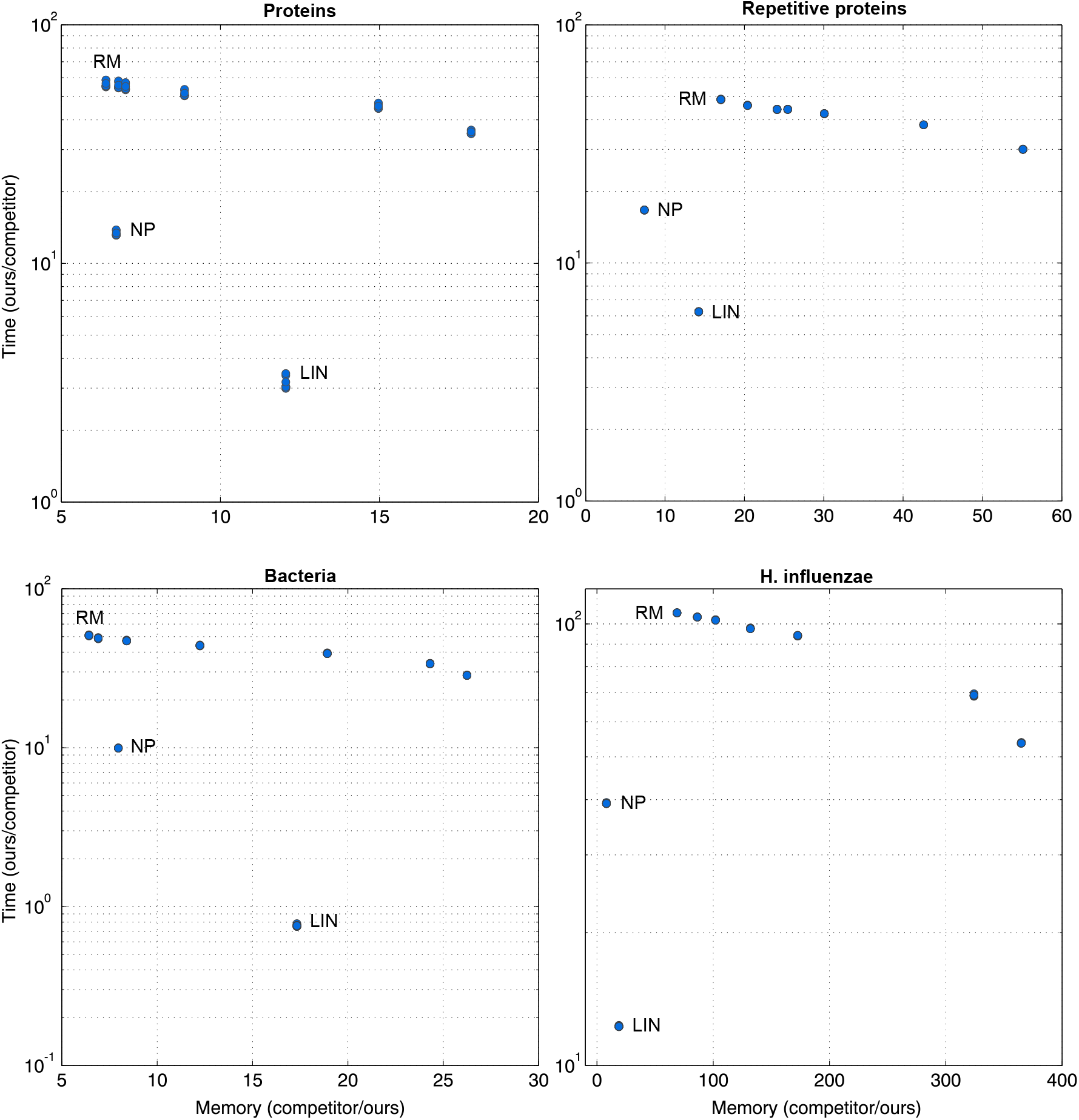
Scoring time: comparison between our index and the implementation in [24]. LIN: score function defined by the competitor. NP: context-based scoring, no pruning. RM: context-based scoring, run-length encoding and maximal repeat pruning. Unlabeled points correspond to RM with decreasing maximum context length (from left to right). Each configuration is tested with ten random substrings, whose measurements are highly overlapping. Memory improvements do not reflect those in Figure 3 of the paper, since the datasets used here are subsets of length at most 80 million of the full datasets.

**Figure 6:**
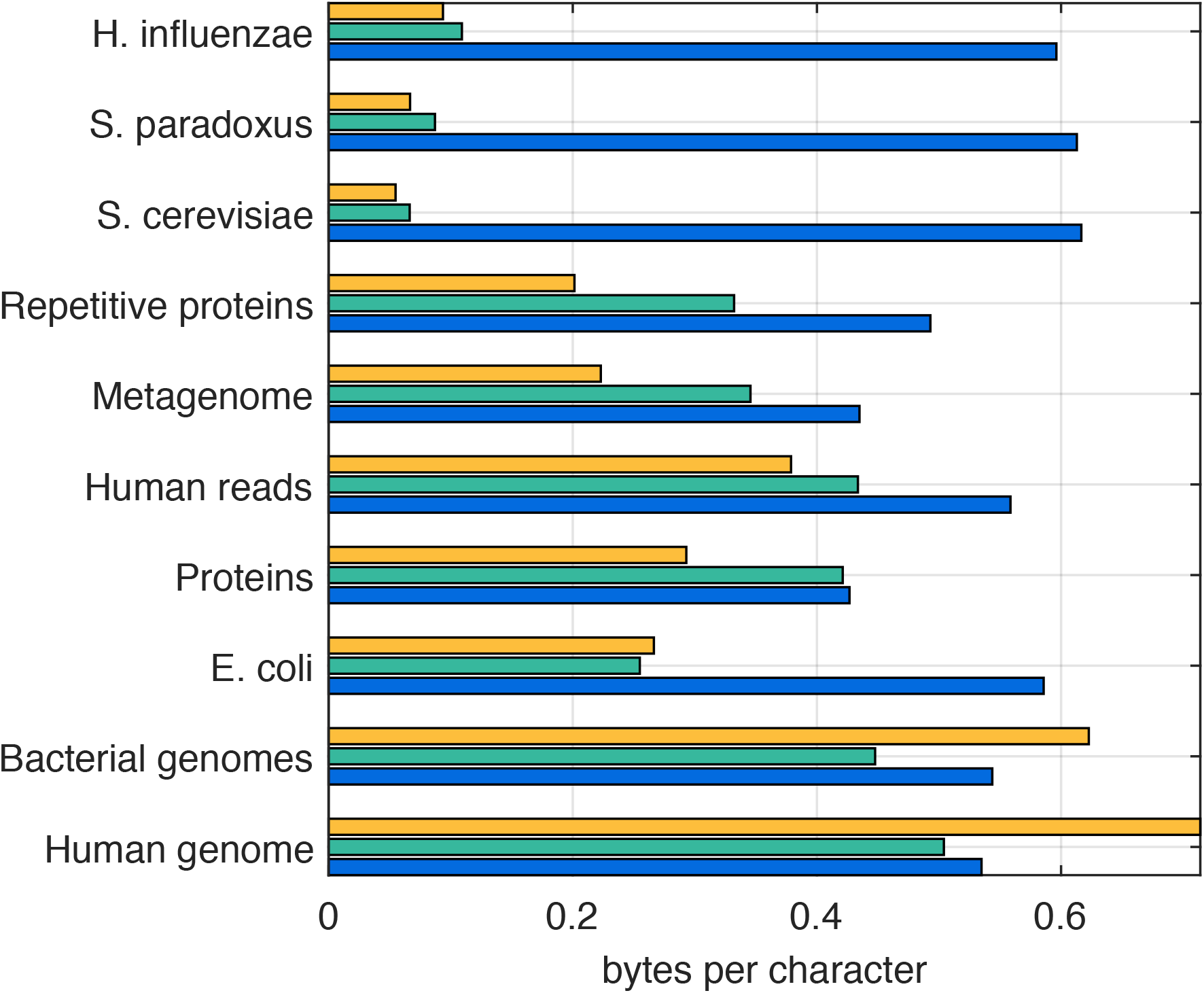
Size of the explicit storage of string depths (orange), compared to the size of the uncompressed (blue) and compressed (green) SLT. Topologies are not pruned.

## Footnotes

1 http://pizzachili.dcc.uchile.cl/repcorpus.html

2 https://emea.illumina.com/platinumgenomes.html

3 https://github.com/eehsan/cstlm

1 https://github.com/y-256/libdivsufsort

2 https://github.com/jnalanko/BDBWTindex

3 https://github.com/nicolaprezza/lz-rlbwt

4 https://www.ncbi.nlm.nih.gov/proteinclusters

5 We observe that setting *b* = 8 achieves a good balance between the size of the array and the number of elements in the hash table.

